# Functional analysis of ligand-gated chloride channels in a cnidarian sheds light on the evolution of inhibitory signalling

**DOI:** 10.1101/2025.07.26.666939

**Authors:** Abhilasha Ojha, Linda Kloss, Alison G. Cole, Audrey Ortega-Ramírez, Juan D. Montenegro, Mihaela Raycheva, Sabrina Kaul, Michèle Bachmann, Sylvia Joussen, Lisa Huf, Simone Albani, Günther Schmalzing, Ulrich Technau, Stefan Gründer

## Abstract

γ-aminobutyric acid (GABA) is the predominant inhibitory transmitter in the vertebrate nervous system. Fast inhibitory signalling is mediated by type A GABA receptors (GABA_A_Rs), which form pentameric ligand-gated ion channels. While GABA is also present in plants and prokaryotes, it is unknown when it was first used for fast neuronal transmission. Cnidaria represent a sister group to all Bilateria and possess a variety of putative GABA_A_Rs, none of which has been functionally characterized. In this study, we surveyed putative inhibitory ion channel receptors from the model cnidarians *Nematostella* and *Hydra*. Phylogenetic analysis revealed a surprising complexity of these receptors. The majority formed a cnidarian-specific radiation with some receptors forming a basal clade. We functionally analyzed seven putative *Nematostella* GABA_A_Rs of this radiation and found that none was activated by GABA or glycine, whereas three were activated by glutamate. Using site-directed mutagenesis, we identified a lysine residue in the canonical ligand-binding pocket that is important for activation by glutamate. Our results identified a group of inhibitory ion channel receptors in Cnidaria that uses glutamate as a ligand. Moreover, they suggest that inhibitory ion channel receptors in Cnidaria massively diversified, which may have been instrumental in the evolution of complex behaviors and sensory processing by the cnidarian nervous system. This work lays the foundation for understanding the diversity and evolution of inhibitory receptors in Cnidaria and the evolution of inhibitory signalling in animal nervous systems.

## Introduction

The balance between excitatory and inhibitory signalling is a hallmark of nervous systems. Fast inhibitory neurotransmission rapidly reduces neuronal activity and is mediated by ligand-gated ion channels (LGICs) that are selective to chloride. The major inhibitory neurotransmitter in the vertebrate nervous system is γ-aminobutyric acid (GABA). LGICs opened by GABA belong to the family of Cys-loop receptors or pentameric LGICs (pLGICs) (Fig.1a). pLGICs include anion-selective and cation-selective members. Cation-selective members include nicotinic acetylcholine receptors (nAChRs) and serotonin 5-HT receptors (5-HTRs). The anion-selective members include GABA_A_ receptors (GABA_A_Rs), glycine receptors (GlyRs), and, in invertebrates, chloride channels gated by glutamate (GluCls) (Cully, et al. 1994; Cully, et al. 1996; Dent 2006), serotonin (MOD-1) (Ranganathan, et al. 2000) and other biogenic amines (Ringstad, et al. 2009), acetylcholine (ACC, LGC-46) (Putrenko, et al. 2005), or histamine (Zheng, et al. 2002). Both GlyRs and GluCls share the unique feature of a second Cys-loop between the signature Cys-loop and the first transmembrane helix M1 (Cully, et al. 1994; Kehoe, et al. 2009). GABA_A_Rs are typically heteropentamers that mediate both synaptic (phasic) and tonic inhibition (Mody and Pearce 2004; Sigel and Steinmann 2012).

Although neurotransmitter signalling through pLGICs is widespread among bilaterians, its evolutionary origin is less clear. Sponges and placozoans do not have nervous systems, whereas in comb jellies (Ctenophora), which are potentially the first diverging animal lineage (Schultz, et al. 2023), only glutamate receptors have been identified (Moroz, et al. 2014). By contrast, in Cnidaria, the sister group to Bilateria, which have diverged about 600 Myr ago, candidates for GABA_A_R can be detected in the genome (Anctil 2009), yet, there is no assessment of the possible biological function of GABA_A_Rs in a cnidarian. A crucial step towards this goal would be the detection of GABA_A_R-expressing cell types and the assessment of their functional properties. A major model organism among cnidarians is the sea anemone *Nematostella vectensis* (Genikhovich and Technau 2009), which has a diffuse nerve net in both cell layers (Nakanishi, et al. 2012; Rentzsch, et al. 2017). Single-cell transcriptome atlases have revealed that *Nematostella* has about 25 distinct neuronal cell types (Steger, et al. 2022; Cole, et al. 2024). Neurogenesis is driven by a conserved set of transcription factors and signalling pathways including SoxC, SoxB2, Achaete-scute and Notch (Layden, et al. 2012; Marlow, et al. 2012; Richards and Rentzsch 2014, 2015; Steger, et al. 2022). This suggests that the neurons are homologous to bilaterian neurons. In addition to neurons, there are also other related neural cell types such as secretory (gland) cells and cnidocytes, the stinging cells (Steinmetz, et al. 2017; Babonis, et al. 2019; Babonis, et al. 2022; Steger, et al. 2022), and muscle of both ectodermal and mesodermal origin (Steger, et al. 2022; Cole, et al. 2023; Cole, et al. 2024). The nervous system is thought to regulate muscle contraction and discharge of cnidocytes; however, it is currently unclear which neurotransmitters might be involved.

Here, we report a large family of putative GABA_A_Rs and GlyRs from *Nematostella vectensis*, most of which belong to a cnidarian-specific radiation of the GABA_A_ subfamily. We cloned and characterized seven pLGICs from this cnidarian-specific radiation and found that none responded to GABA or glycine, but three were activated by glutamate. We identified a unique lysine residue in the ligand-binding pocket, which is essential for glutamate sensitivity. Our study is the first to functionally characterize recombinant pLGICs from a cnidarian species and indicates that Cnidaria use glutamate and probably other transmitters besides GABA and glycine for fast inhibitory transmission via pLGICs. Our results lay the foundation for a better understanding of the evolution of fast inhibitory signalling in Cnidaria and in animal nervous systems in general.

## Results

### Most putative GABA_A_Rs in *Nematostella* belong to a cnidarian-specific radiation and some form homopentamers

In the newly annotated transcriptome that accompanies the chromosome-level genome assembly of *Nematostella vectensis* (Zimmermann, et al. 2023), we identified a total of 42 pLGICs and 15 pLGICs from *Hydra vulgaris* (strain AEP) (Cazet, et al. 2023) with homology to the GABA_A_R/GlyR superfamily, which were included in thorough phylogenetic analyses with other bilaterian pLGICs. We performed maximum likelihood (ML) and Bayesian analyses, using mammalian 5-HTRs as an outgroup, with essentially similar results (Fig. 1 and Supplementary Fig. S1). The phylogenetic analyses showed that of the 42 *Nematostella* receptors, most (37 in the ML analysis shown in Fig. 1) belong to a cnidarian radiation of putative GABA_A_Rs, which also includes 10 receptors from *Hydra*. Within this cnidarian-specific radiation, in the ML analysis, there are eight clades, two with representatives from *Hydra*, indicating a higher diversification of inhibitory receptors in Anthozoa than in Hydrozoa. In The Bayesian analysis, the relation of cnidarian-specific clades I and II to the other clades was not resolved and, therefore, remains ambiguous. In both analyses, 3 *Nematostella* receptors (NV2.24331, NV2.3907, and NV2.13369) are placed more closely to vertebrate GABA_A_Rs, however, the support values are low. Notably, 2 other *Nematostella* receptors (Nv2.17304 and Nv2.15844) robustly cluster together with 2 *Hydra* proteins at the base of the GABA_A_R/GlyR superfamily with good support from ML analysis (bootstrap 84 %) and modest support from Bayesian inference (posterior probability 0.74). As a conclusion, our analyses reveal a massive cnidarian expansion of LGICs, in particular in *Nematostella*, as a sister to the bilaterian GABA_A_R/GlyR superfamily. While several internal clusters are well-supported, the evolutionary relationship of the clusters with each other cannot be reconstructed. The robust position of a small cluster of 2 *Hydra* and 2 *Nematostella* receptors at the base of cnidarian/bilaterian GABA_A_R/GlyR superfamily suggests that these have retained more ancestral features. Because of this complex phylogenetic history, automated gene annotations based on best BLAST hits should be taken with caution. Notably, several experimentally validated acetylcholine- and serotonin-gated channels from *C. elegans* (ACC, LGC-46, MOD-1) seem to be the channels most closely related to the cnidarian-radiation. This indicates that the ligand specificity cannot be easily predicted from the phylogenetic analysis alone.

**Figure 1.**
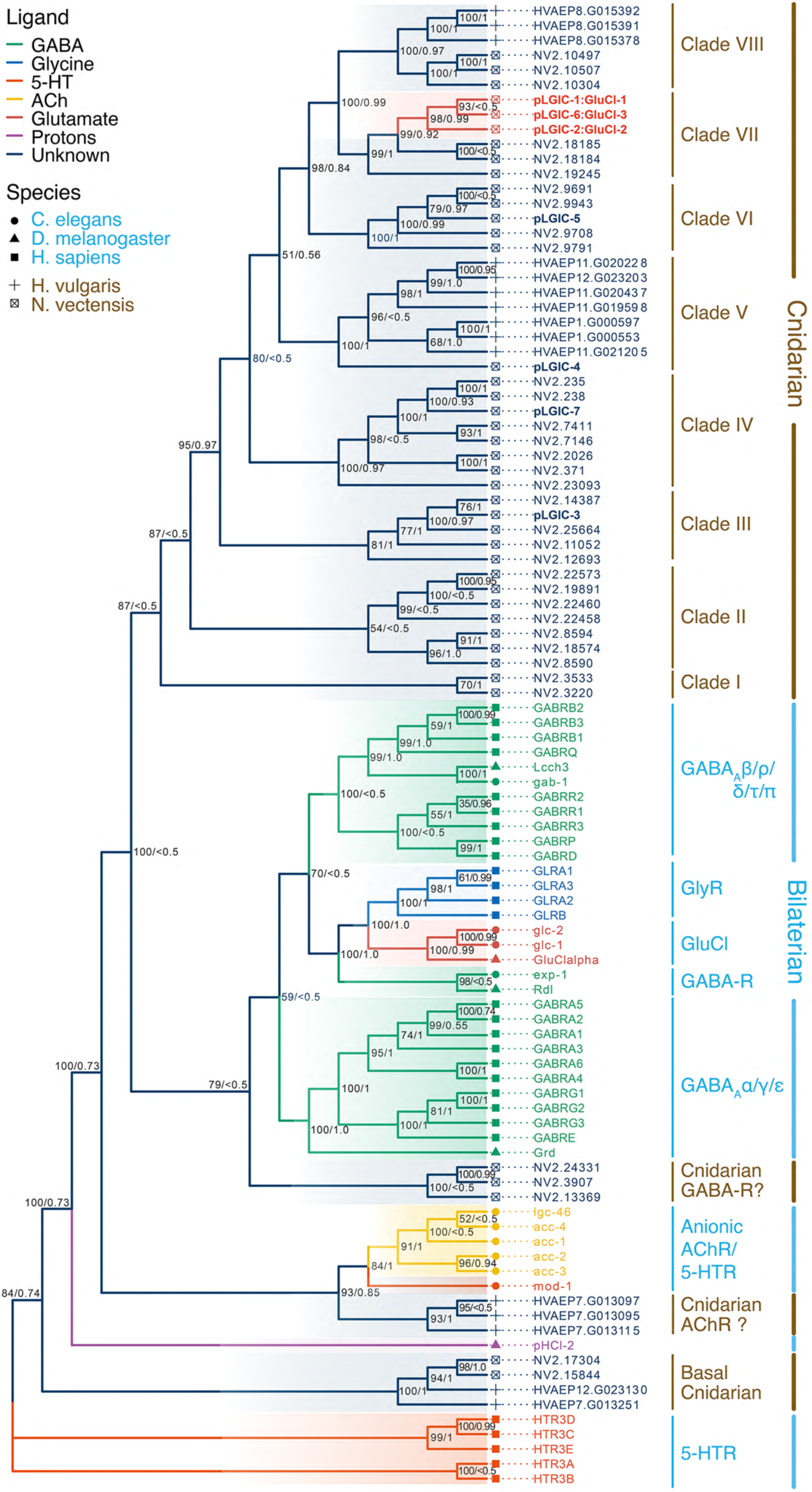
Phylogenetic analysis of cnidarian GABA_A_Rs/GlyRs. Maximum likelihood analysis of pLGIC genes in *N. vectensis*, *H. vulgaris*, *C. elegans*, *D. melanogaster*, and *H. sapiens*. The cnidarian-specific expansion of GABA_A_R/Gly-like receptors contains eight clades. Multispecies GABA_A_/Gly receptors branch basally to these clades, suggesting a cnidarian-specific radiation. Cloned receptors of this study are shown in bold. Note that the relation of bilaterian GABA_A_-α, GABA_A_-β and GlyRs to each other has low support. Annotation of three receptors from *N. vectensis* as “Cnidarian GABA-Rs” and of three from *H. vulgaris* as “Cnidarian AChR” is provisional as it has relatively low support. Support values for individual nodes are indicated (bootstrap/posterior probability). The tree was rooted with mammalian 5-HT_3_Rs as an outgroup.

For further characterization, we cloned seven subunits from *N. vectensis*, referred to here as nvpLGIC-1 –nvpLGIC-7 (red in Figure 1), belonging to five of the eight gene clades of the cnidarian-specific radiation. nvpLGIC-1 – nvpLGIC-7 are 405 – 490 amino acids in length (Supplementary Figure 2). Sequence alignment with the human GABA_A_ α1 and β3 subunits revealed that they contain the hallmarks of Cys-loop receptors: an N-terminal ligand-binding extracellular domain (ECD), four transmembrane helices (M1-M4), an intracellular loop between M3 and M4, an extracellular C-terminal domain, and a disulfide bond in a loop at the interface between the ECD and the transmembrane domain, which identifies them as Cys-loop receptors (Fig. 2a and Supplementary Fig. S2). Human α1 GABA_A_ subunits form homopentamers when a residue in either M1 or M3 is mutated to Trp (Hannan and Smart 2018), and an intramembrane aromatic network determines the assembly of pLGICs (Haeger, et al. 2010). nvpLGIC-1 – nvpLGIC-7 contain a Trp residue at the critical position in M1 (Supplementary Fig. S2), suggesting that they might form homopentamers. To directly test the assembly and formation of homopentamers, we tagged three representative pLGICs (pLGIC-1, pLGIC-3, and pLGIC-5) at their extracellular C-termini with a hexahistidine-tag and analysed their plasma membrane expression and oligomeric assembly state using denaturing SDS-PAGE and non-denaturing blue native PAGE after expression in *Xenopus laevis* oocytes. As a positive control, we used the human GABA_A_ β3 subunit, hGABRB3, which assembles as a functional homopentamer, the structure of which has been solved (Miller and Aricescu 2014). We purified the histidine-tagged versions of these receptors under non-denaturing conditions using Ni-NTA chromatography. The total pool of GABA subunits was metabolically labelled with [^35^S]methionine, whereas the plasma membrane-bound pool was labelled at the cell surface of intact oocytes by the lysine-reactive infrared dye IR800-NHS ester. The purified nvGABA receptors migrated both in the plasma membrane form (Fig. 2b) and in the whole cell form as distinct high-mass proteins that dissociated into lower-order intermediates when exposed to the denaturing detergent SDS for 1 h at 37°C (Fig. 2b). The maximum number of five bands generated indicates that all three nvGABA receptors assemble as homopentamers, identical to hGABRB3. The similarity of the band patterns reflects the similar subunit masses of the four proteins. Using the crystal structure of the human β3 homopentamer (Miller and Aricescu 2014) as a template, we generated a homology model of homomeric nvpLGIC-2, revealing a canonical pLGIC structure with a Cys-loop and several loops that form the ligand-binding site (Fig. 2c).

**Fig. 2.**
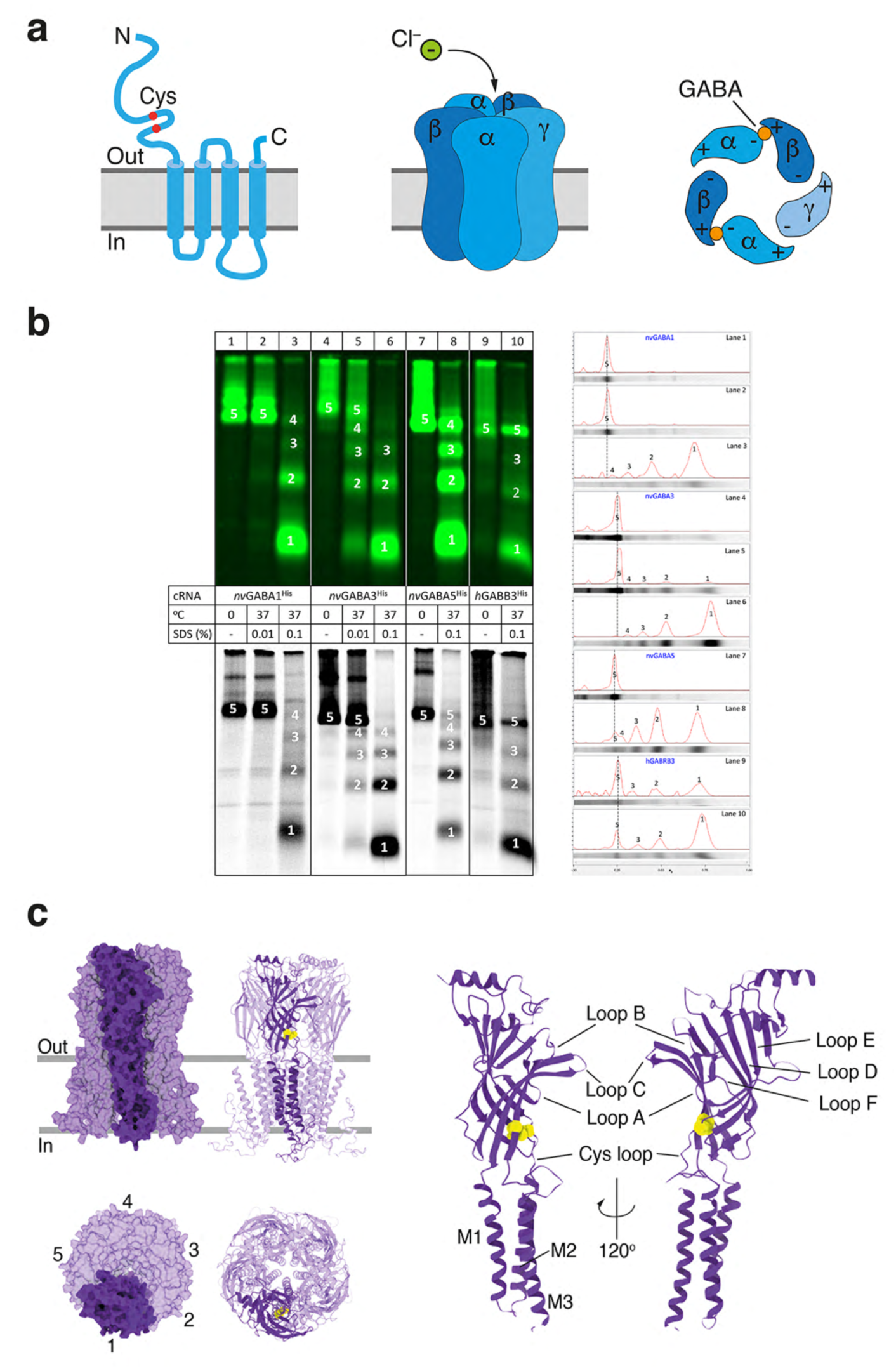
*Nematostella* pLGICs form homomeric Cys-loop receptors. **a)** Schematic representation of GABA_A_Rs. Left, cartoon illustrating the transmembrane topology of a single subunit. The characteristic Cys-loop is indicated. Middle, cartoon illustrating the pentameric structure. In mammals, typical GABA_A_Rs have a 2α2βγ stoichiometry. Right, top-view cartoon, illustrating the binding of two GABA molecules at the β+/α– interfaces. **b)** Visualization of the oligomeric state of nvpLGICs versus pentameric human GABA_A_ β3. Top left, IR800 fluorescence detecting the surface-expressed pool of receptors. Bottom left, ^35^S-labelled proteins detecting the total pool. Right, IR800 fluorescence was quantified along each lane, which is also shown horizontally in grey. Peaks are numbered according to the protein bands on the left. The close similarity of the band patterns to the similarly sized human β3 receptor identifies the nvpLGICs as homopentamers. **c)** Homology model of nvpLGIC-2 based on PDB entry 4COF. Left, overview of the homopentameric complex. A single subunit is highlighted. Top, side view; bottom, top view. Right, domain structure of a single subunit. S-S bonds are shown as yellow spheres. Loops contributing to the ligand binding site are indicated. Presentation based on Allard et al. (Allard, et al. 2023).

### Functional expression revealed three glutamate-gated Cl^-^-channels (nvGluCls) among putative GABA_A_ receptors

We expressed nvpLGIC-1 – nvpLGIC-7 individually in *Xenopus* oocytes and functionally characterized them using two-electrode voltage clamp. Surprisingly, neither GABA nor glycine (1 μM – 1 mM) evoked currents in nvpLGIC-expressing oocytes. We then considered glutamate, the precursor of GABA, as a putative small-molecule agonist. Interestingly, glutamate evoked large currents in oocytes expressing nvpLGIC-1, nvpLGIC-2, and nvpLGIC-6, but not in oocytes expressing nvpLGIC-3, nvpLGIC-4, nvpLGIC-5, nvpLGIC-7 (Fig. 3a). Similarly, serotonin, D-serine, and acetylcholine did not elicit currents with nvpLGIC-1 – nvpLGIC7 (Fig. 3a) nor did aspartate, glutamine, histamine, or β-alanine, not even with a mix of nvpLGIC-1 – nvpLGIC-7 (we did not test nvpLGIC-7 individually with the latter agonists). Therefore, we renamed the three receptors responding to glutamate nvGluCl-1 (nvpLGIC-1), nvGluCl-2 (nvpLGIC-2), and nvGluCl-3 (nvpLGIC-6), respectively. These three receptors form a monophyletic clade within the cnidarian GABA_A_ family (Fig. 1).

**Fig. 3.**
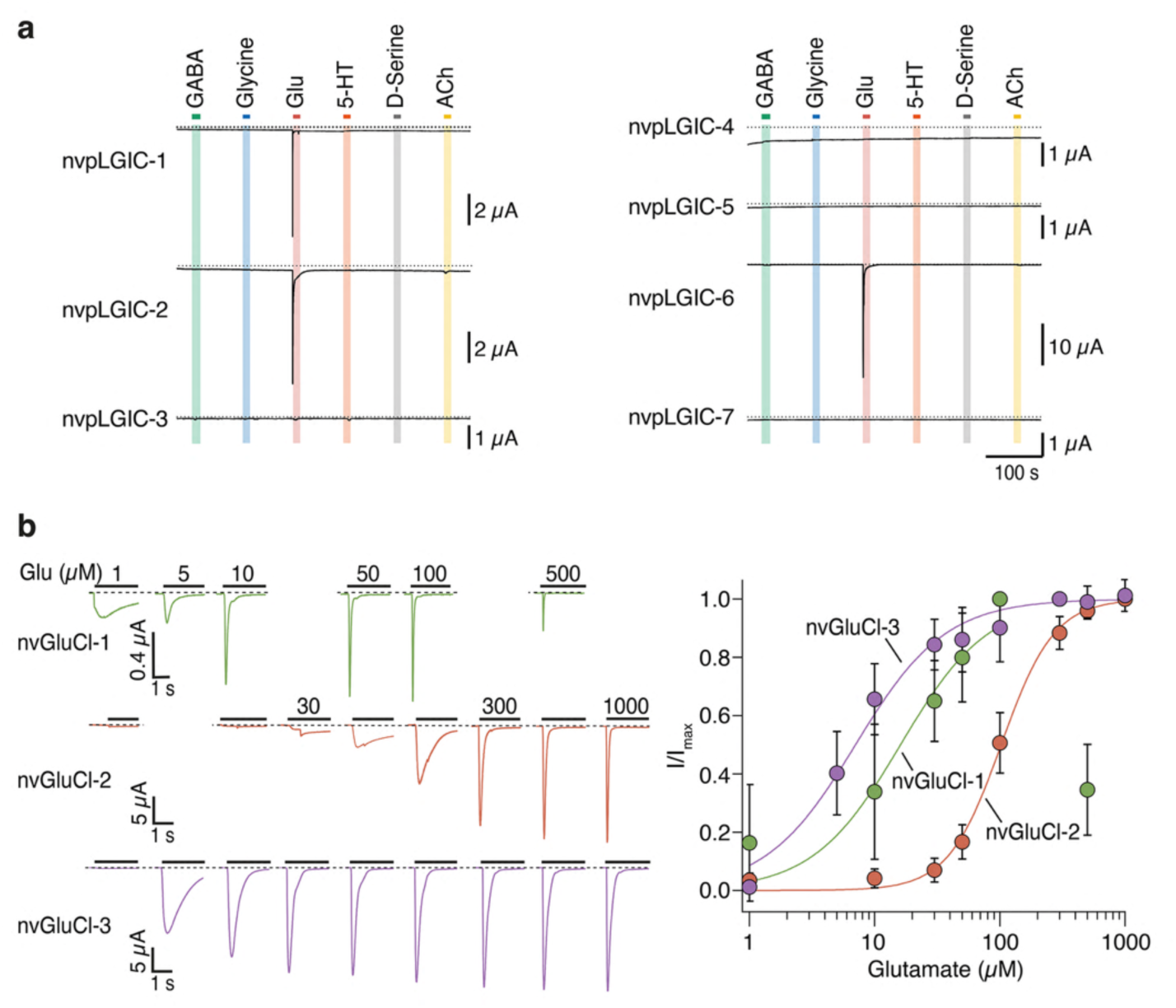
Three *Nematostella* pLGICs are activated by micromolar concentrations of glutamate. **a)** representative current traces from oocytes expressing nvpLGIC-1 to nvpLGIC-7. The indicated ligands were applied at a concentration of 1 mM, except for glutamate, which was applied at 100 μM. Dashed lines represent the 0 current level. **b)** Left, representative current traces showing activation of nvGluCl-1 (nvpLGIC-1), nvGluCl-2 (nvpLGIC-2), and nvGluCl-3 (nvpLGIC-6) upon application of the indicated concentrations of glutamate (black bars). Right, concentration-response curves for nvGluCl-1 (green), nvGluCl-2 (orange), and nvGluCl-3 (violet). Data represent mean normalized currents ± s.d. from 6-17 cells. Solid lines indicate fits to the Hill function.

Application of increasing glutamate concentrations revealed that half-maximal activation of the three receptors varied more than 10-fold, ranging from 7 to 100 µM (EC_50_ = 16 µM for GluCl-1, EC_50_ = 102 µM for GluCl-2, and EC_50_ = 7 µM for GluCl-3, respectively; Fig. 3b). Glutamate-evoked currents desensitized completely. With increasing glutamate concentrations, desensitization became more rapid until it reached a plateau at saturating glutamate concentrations. We determined the time constant of desensitization, τ_des_, at 500 μM glutamate with a fit to a single exponential function, revealing τ_des_ = 26 ± 3 ms for nvGluCl-1, 111 ± 47 ms for nvGluCl-2, and 161 ± 93 ms for nvGluCl-3, respectively (Supplementary Fig. 3a). It is likely that these time constants are limited by the comparatively slow solution exchange (10-90 % solution exchange ∼300 ms (Chen, et al. 2006)) of our setup for *Xenopus* oocytes. However, these values are similar to those of mammalian GABA_A_Rs, which typically desensitize with a fast component on the order of 2-50 ms and a slow component on the order of 50 ms to several seconds, depending on the subunit stoichiometry (Jones and Westbrook 1995; Berger, et al. 1998; Papke, et al. 2011). For nvGluCl-2 and nvGluCl-3, we also determined recovery from desensitization, which had a similar τ_recovery_ of 31 sec and 38 sec, respectively (Supplementary Fig. 3b).

To assess the ion selectivity of nvGluCl-2 and nvGluCl-3, we determined the reversal potential E_rev_ with 140 mM NaCl in the extracellular solution, revealing an E_rev_ of ∼-20 mV, close to the reversal potential of Cl^-^ in oocytes under these ionic conditions. When Cl^-^ in the extracellular solution was replaced with the large impermeant anion gluconate, E_rev_ shifted to ∼+30 mV (Fig. 4a), confirming that Cl^-^ was the main permeant ion. The ion pore of Cys-loop receptors is lined by residues of the five M2 α helices. Figure 4b shows an alignment of the M2 sequences of nvGluCl-1, nvGluCl-2, and nvGluCl-3, together with human α1 and β3 GABA_A_ subunits. The open ion pore of pLGICs has its strongest constriction at the -2′ residue (Hibbs and Gouaux 2011; Miller and Aricescu 2014). Determinants of anion selectivity are -1′Ala and -2′ Pro residues, whereas cation channels have a -1′ Glu residue (Hibbs and Gouaux 2011). In nvGluCl-1-nvGluCl-3, 1′Ala and -2′ Pro are conserved, consistent with the finding that they are glutamate-gated Cl^-^ channels. In addition, the homology model of nvGluCl-2 confirmed the constriction at Pro-2′ and the position of residues lining the ion pore (Fig. 4c).

**Fig. 4.**
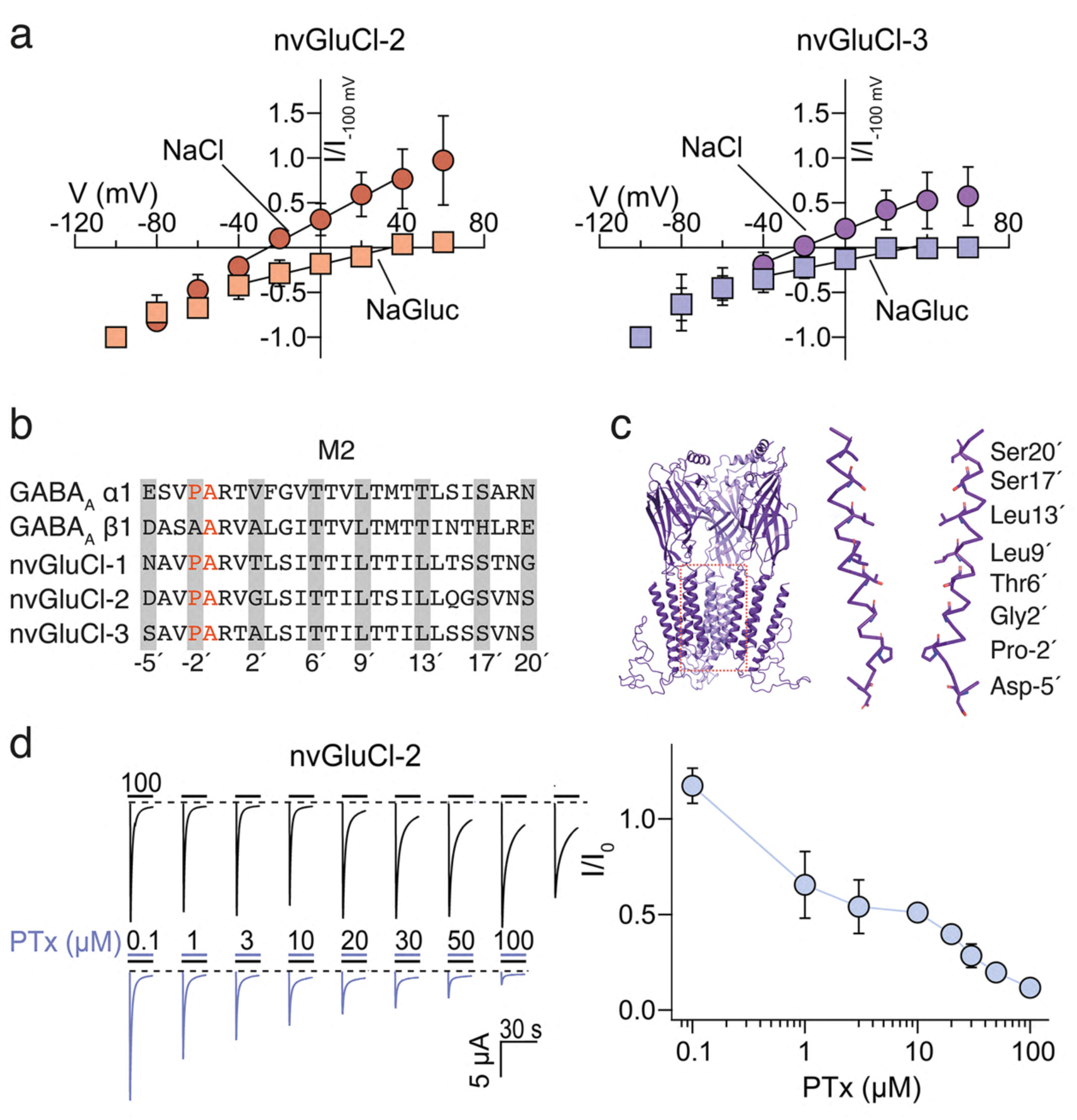
nvGluCls are chloride channels. **a)** I-V relationships for nvGluCl-2 (left) and nvGluCl-6 (right) in control bath solution (circles) and in a bath solution, in which NaCl was replaced by NaGluconate (NaGluc; squares). nvGluCl-2 was activated with 500 µM glutamate, and nvGluCl-6 (right panel) with 50 µM glutamate. Data are plotted as mean ± s.d. for 3-15 cells. Data points were fitted with a linear function (black lines) to obtain the reversal potential (*E*_rev_). For nvGluCl-2, *E*_rev_ was -25 mV in control solution and 34 mv with NaGluconate, and for nvGluCl-6, *E*_rev_ was -21 mV in control solution and 29 mV with NaGluconate. **b)** Sequence alignment of M2 sequences of human GABA_A_ α1, human GABA_A_ β3, and nvGluCl-1-nvGluCl-3. The cytoplasmic side is to the left, and the extracellular side is to the right. Residues that line the ion pore are shown on a grey background and numbered according to the convention for Cys loop receptors. They form rings of identical residues in five-fold symmetry of functional homomeric receptors. The-1′Ala and -2′ Pro residues are shown in red. **c)** Homology model of nvGluCl2. The front subunit has been removed for clarity, and the indicated region is shown on an expanded scale on the right. Only two M2 helices are shown. Amino acids facing the ion pore are shown as sticks. **d)** Left, representative current traces for nvGluCl-2 obtained upon application of 100 µM glutamate with and without the indicated concentrations of picrotoxin (PTx). Right, concentration-response curve. Data represent the mean ± s.d. of 6-16 cells.

To further test the conservation of the nvGluCl ion pore with the ion pores of GABA_A_Rs, we assessed the inhibition of nvGluCl-2 by picrotoxin (PTx), an open-channel blocker of GABA_A_Rs. PTx binds to the cytosolic base of the pore, close to M2 residues -2′and 2′ in *Caenorhabditis* GluClα (Hibbs and Gouaux 2011) or between the M2 2′ and 9′rings in the heteromeric human α1β3γ2L GABA_A_R (Masiulis, et al. 2019). In the human receptor, the hydrophobic isoprenyl moiety of PTx is surrounded by the 9′ Leu ring, and the oxygen atoms of PTx form putative hydrogen bonds with the 6′ Thr ring. The conservation of the 6′ Thr and 9′ Leu rings in nvGluCls (Fig. 4b and 4c) strongly suggests that they could also bind PTx. PTx indeed inhibited nvGluCl-2 with an apparent IC_50_ of ∼10 μM (Fig. 4d), similar to canonical GABA_A_Rs, illustrating the conserved overall architecture of the nvGluCl ion pore with that of GABA_A_Rs.

### Amino acid residues that determine the ligand sensitivity of nvGluCls

We next addressed the question of which specific amino acids in the ligand-binding pocket are responsible for glutamate versus GABA binding in nvGluRs. The ligand-binding pocket of Cys-loop receptors localizes at the interface of two neighbouring subunits. One subunit contributes the principal or (+) side, and the other the complementary or (-) side (Fig. 2a). The binding pocket is mainly formed by non-contiguous loops A-C on the (+) side and two β-strands and one loop (“loops” D-F) on the (-) side (Fig. 2c, 5a) (Lynagh and Pless 2014). Human α1β2γ2 and α1β3γ2 GABA_A_Rs contain two α subunits and two β subunits and the GABA-binding pockets are located at the two β-α interfaces where the β subunits contribute the (+) sides (Fig. 2a). GABA is surrounded by an “aromatic box” formed by Y97, Y157, F200, and Y205 of the β subunit, and F65 of the α subunit. While the GABA amino group is engaged in a cation-τ interaction with β-Y205 of loop C and is further stabilized by hydrogen bonds with β-E155 of loop B, the γ-carboxylate of GABA forms a salt bridge with α-R67 of loop D (Zhu, et al. 2018; Masiulis, et al. 2019). Overall, key amino acids of the ligand-binding pocket are surprisingly well conserved in the three nvGluCls (Fig. 5a). In particular, the Arg residue of loop D and the aromatic box are completely conserved, except for the residue corresponding to β-F200 of loop C, which is replaced by a polar Asn residue. Moreover, nvGluCls have a Lys residue in loop C (K229 in nvGluCl-2; Fig. 5a), close to the site where the basic GABA amino group binds in human GABA_A_Rs. This lysine should increase the positive electrostatic potential of the binding pocket and might help to stabilize the additional α-carboxylate group of the glutamate ligand. The homology model of nvGluCl-2 confirmed that loops A-F could form a classical neurotransmitter binding site (Fig. 2c). To obtain a better impression of the potential Glu-binding site in nvGluCls, we docked Glu to the nvGluCl-2 homology model (Fig. 5b). Strikingly, the best binding pose indicated a salt bridge between the α-carboxylate of Glu and K229 of loop C. In this model, R67 of loop D forms a salt bridge with the α-carboxylate of Glu, instead of the γ-carboxylate of the ligand as in GABA_A_Rs. In the nvGluCl-2 model, the γ-carboxylate of Glu rather forms a salt bridge with K74 of a different region (“loop G”) from the (-) subunit. Interestingly, in GluClα of *C. elegans*, an Arg residue of loop G (Fig. 5a) interacts with the α-carboxylates of glutamate (Hibbs and Gouaux 2011). Thus, it appears that the glutamate ligand is anchored in the ligand-binding pocket of nvGluCl-2 via electrostatic interactions between its two carboxylates and two Lys residues (Fig. 5b). To test the contribution of K229 to ligand binding in nvGluCl-2, we mutated it to threonine, the residue found in human β subunits, and found that this mutation completely abolished the response to high concentrations of glutamate (Fig. 5c), confirming that K229 makes an essential contribution to glutamate binding. Mutant channels were still not activated by GABA (Fig. 5c), indicating that other residues are also important for the ligand switch. Importantly, a Lys residue is present in loops C and G only of nvGluCl-1 – nvGluCl3, but not in nvpLGIC-3, nvpLGIC-4, nvpLGIC-5, or nvpLGIC-7 (Supplementary Fig. 2), further suggesting that these residues are important for glutamate binding in nvGluCls and that nvpLGICs lacking these Lys residues have another ligand.

**Fig. 5.**
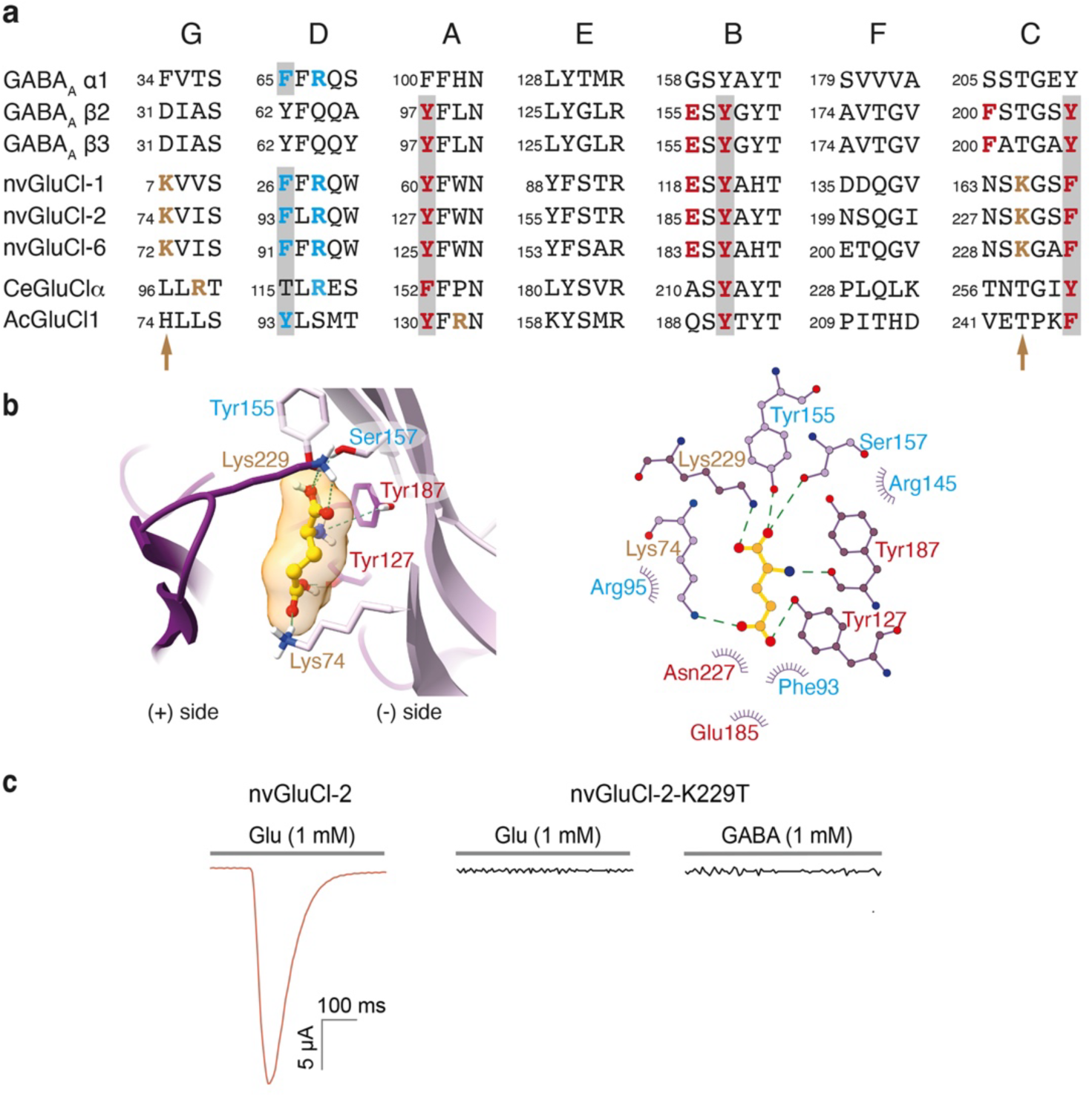
A lysine residue in loop C is essential for glutamate binding. **a)** Sequence alignments of the loops forming ligand-binding pockets of human α1, β2, and β3 subunits, of nvGluCls and GluCls from *Caenorhabditis elegans* and *Aplysia californica*. Crucial residues of the principal β subunits are shown in red and those of the complementary α subunits are shown in blue; residues of the aromatic box are on grey background. The critical Lys residues in loops C and G of nvGluCls are shown in brown (arrows). **b)** Result from docking Glu onto nvGluCl-2. Left, close-up of the binding pocket revealing electrostatic interactions of Glu with K74 and K229. Right, LIGPLOT schematic of Glu interactions. Electrostatic interactions are shown as dashes and hydrophobic interactions as eyelashes. Residues of the principal and complementary subunits are shown in red and blue, respectively. K74 and K229 are shown in brown; K74 is from the complementary subunit, and K229 is from the principal subunit. **c)** Loss of function of nvGluCl-2 with the K229T mutation.

### Cnidarian-specific pLGIC receptors are expressed in cnidarian-specific cell types and neurons

We assayed the expression profiles of the nvpLGICs in the available single-cell atlas (Cole, et al. 2024) and found that 36 of them are specifically expressed at detectable levels across various cell states (Supplementary Fig. S4). Expression was predominantly found within mature cnidocyte profiles and related N2 class neurons, defined by the expression of Brn3/POU4 transcription factor family gene (Cole, et al. 2024). nvGluCls were detected within cell types specific to the tentacles of the polyp, including both spirocytes, an anthozoan-specific cell type (Sierra and Gold 2024), and tentacle retractor muscles (Fig. 6a). nvGluCl-3 was also detected in one specific neural subtype (N2 neuron 3), whereas pLGIC-5 is expressed in both mature spirocytes and the FoxQ2d-positive N1S neuron 4 (Busengdal and Rentzsch 2017; Cole, et al. 2024). To validate the predicted gene expression from single-cell RNA-seq analyses, we generated in situ hybridization probes for the gene models of the three GluCl-responsive receptors. We found expression of all three genes from the late planula stage, which accumulate in the tentacle bud stage in the growing tentacles during metamorphosis into the primary polyps (Fig. 6b-d). Once the tentacles have elongated, nvGluCl-1 and nvGluCl-3 are expressed in individual cells at the base of the ectoderm (Fig. 6g), which is indicative of expression in tentacle neurons and tentacle retractor muscle cells. In addition, nvGluCl-1 and nvGluCl-2 are expressed in ectodermal cells at the tentacle tip (Fig. 6e and 6f). This spatial distribution is characteristic of spirocytes, a type of cnidocyte that is restricted to the tentacles.

**Fig. 6.**
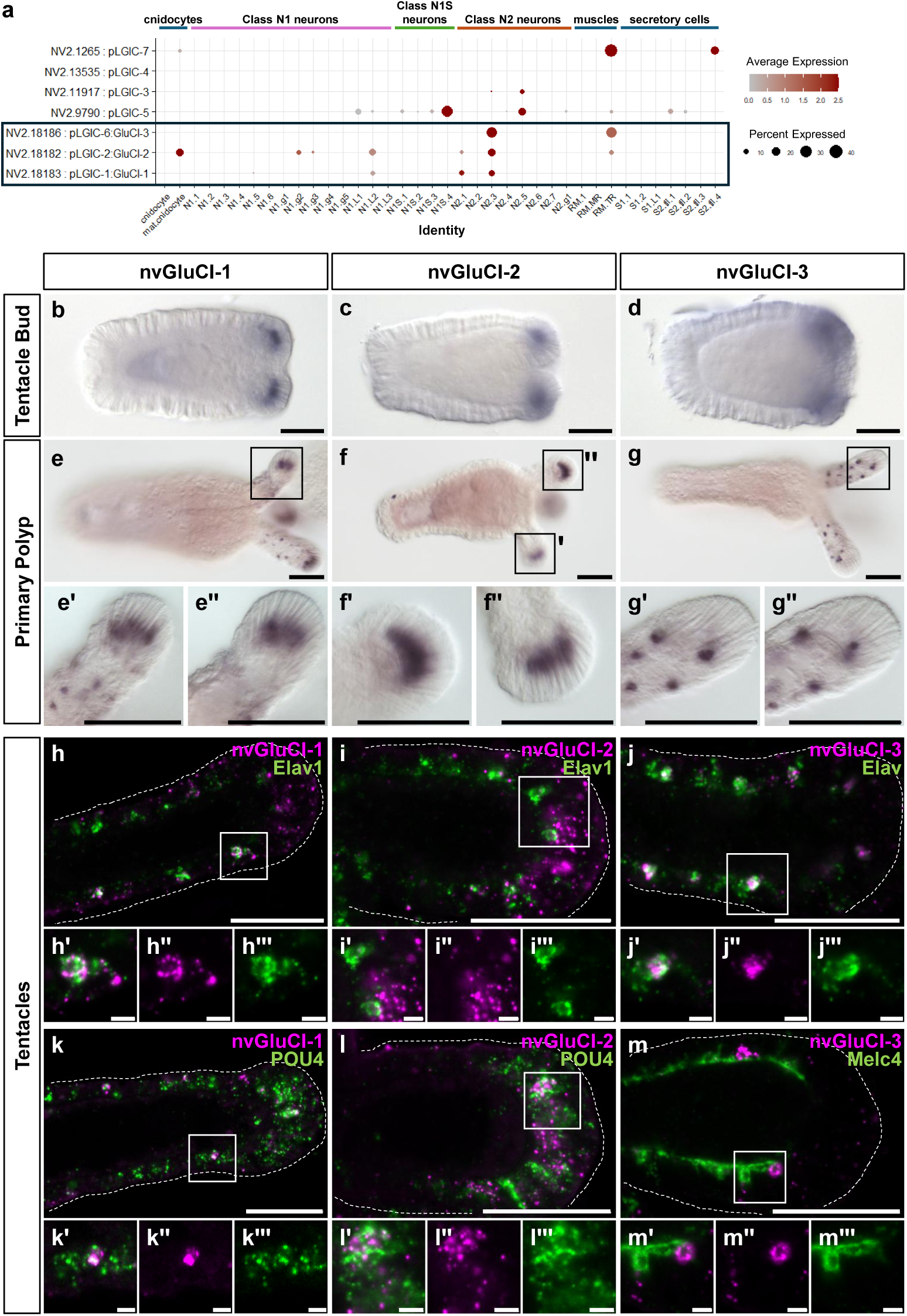
pLGIC expression is predominately in cells of the tentacle and individual class N2 neurons. **a)** Dotplot of expression profiles within cnidocytes, neurons, secretory cells, and fast muscle of the seven candidate receptors investigated in this study. Expression in early cell states is not shown. See supplementary figure S2 for expression profiles of all genes across the entire dataset. The three highlighted genes were further investigated using *in situ* hybridization. **b to g)** In situ hybridization showed expression within cells of the tentacles from the tentacle bud to the primary polyp stage (indicated on the left). All images show lateral views with the aboral pole to the left. Scale bars, 50 µm overview, 10 µm enlarged section. **h–m)** Double fluorescent *in situ* hybridization for GluCl (magenta) and cell type-specific markers (green) confirmed expression of nvGluCl-1 in neurons (h, k), nvGluCl-2 in spirocytes (i, l), and nvGluCl-3 in cells closely associated with the tentacle retractor muscle (j, m). Images are confocal sections showing lateral-view tentacles of primary polyps with tentacle tips to the right. Scale bars, 50 µm overview, 5 µm enlarged section.

To confirm the cell types in which the three receptors are expressed, we performed double fluorescence *in situ* hybridization (Fig. 6h and 6i). This showed that nvGluCl-1 and nvGluCl-3 are specifically expressed in a subpopulation of Elav1-positive neurons (Nakanishi, et al. 2012) (Fig. 6h). Furthermore, nvGluCl-1 and nvGluCl-2 are expressed in POU4-positive cells (Tourniere, et al. 2020; Steger, et al. 2022), which potentially include both N2 neurons and cnidocytes (Fig. 6k and 6l), in line with the idea that these neurotransmitter receptors define the mature state of the cnidocytes. We also investigated the expression of nvGluCl-3 in the tentacle retractor muscle by double fluorescent *in situ* hybridization with Melc4 as a marker of fast-retracting muscle (Cole, et al. 2023) (Fig. 6m). nvGluCl-3 seems to be expressed in cells closely associated with tentacle retractor muscle cells, presumably neurons innervating the tentacle retractor muscles. Occasional co-expression with Melc4 cannot be ruled out because of its close co-localization (Fig. 6m). Together, these observations suggest that the three tested nvGluCl-responsive receptors may be involved in the coordination of tentacle movement in response to predators or prey.

## Discussion

### The evolution of inhibitory signalling

Neuronal LGICs are fundamental for complex behaviours and rapid sensory processing. However, the evolution of neurotransmitter signalling through LGICs in animals remains an unsolved mystery. Ionotropic glutamate receptors (iGluRs) have an ancient evolutionary origin, as they are also present in plants (Lam, et al. 1998) and likely evolved from prokaryotic ion channels (Chen, et al. 1999). The origin and ancient ligands of pLGICs are less clear. While sponges have iGluRs but lack AChRs and GABA_A_Rs, cnidarians, the sister group to Bilateria, contain divergent copies of genes coding for iGluRs and of genes with homology to AChRs, GABA_A_Rs, or GlyRs (Anctil 2009). Our results reveal a surprising complexity of the genes with homology to GABA_A_Rs/GlyRs in the cnidarian *Nematostella*. Of the 42 nvpLGICs that belong to the inhibitory pLGIC branch, two are forming a basally branching cluster with two Hydra pLGICs. In the ML analysis, 37 *Nematostella* and 10 *Hydra* LGICs form an impressive cnidarian expansion with a likely sister relationship to the bilaterian receptors. These findings have important implications for the evolution of inhibitory LGICs.

First, it remains unclear whether the ancestor of vertebrate GABA_A_Rs/GlyRs existed before the cnidarian-bilaterian split. While our phylogenetic analyses put three *Nematostella* receptors on the same clade as vertebrate GABA_A_Rs/GlyRs, this position has relatively low support and therefore remains uncertain. However, the specialization of vertebrate subunits, for example into α and β subunits, likely occurred after this split. More importantly, whether the ancestor of vertebrate and cnidarian pLGICs was gated by GABA remains an open question. GABA is an important inhibitory neurotransmitter in both vertebrates and invertebrates. While prokaryotic cells and plants can also synthesize GABA, where it serves as a metabolic product, the important question is when GABA was first used for neuronal signalling in animal nervous systems. *N. vectensis* possesses genes with homology to the key enzyme for GABA synthesis, glutamate decarboxylase (GAD), with a relatively low identity to the vertebrate enzymes (Anctil 2009; Oren, et al. 2014). While their expression seems to be restricted to an endodermal cell layer around the pharynx and testis, but absent from the tentacles (Marlow, et al. 2009), GABA immunoreactivity has been found in sensory and ganglion cells and in cnidocytes of *N. vectensis* (Marlow, et al. 2009). *N. vectensis* also has homologs of proteins required for GABA import into synaptic vesicles, presynaptic reuptake, and metabotropic GABA_B_ receptors (GABA_B_Rs), which contain conserved GABA-binding sites and presumably mediate the effects of the GABA_B_-specific agonist baclofen on neurogenesis (Levy, et al. 2021). Thus, metabotropic GABA_B_Rs likely mediate GABA signalling in *N. vectensis*. Homology-based prediction of GABA_A_Rs in the *N. vectensis* genome led to the assumption that *N. vectensis* uses GABA also for fast ionotropic signalling. In TEVC experiments, of seven expressed LGICs, none was activated by GABA or glycine, but three were shown to be chloride channels activated by glutamate. Since all three are expressed in cnidocytes, tentacle rectractor muscle or neurons in the tentacle, we speculate that the receptors might be involved in the control of the feeding response. Of note, among the bilaterian pLGICs of the GABA_A_R/GlyR superfamily, there are also several AChRs from *C. elegans*, suggesting the independent acquisition of this ligand in this animal lineage. Taken together, the combination of phylogenetic and functional analyses used in our study suggests that the radiation of LGICs preceded the acquisition of ligand specificity, which evolved independently within the various clades and animal lineages. In summary, the answer to the question of the ancient ligand of channels of the GABA_A_R/GlyR superfamily remains open and awaits deorphanization of more cnidarian receptors within the GABA_A_R clade (e.g. NV2.24331, NV2.3907, and NV2.13369) and of those basal to all other inhibitory pLGICs (NV2.17304 and NV2.15844).

### The diversity of inhibitory LGICs in *Nematostella*

The second implication of our findings is that inhibitory pLGICs diversified extensively in cnidarians. In the ML and Bayesian analyses, it appears that all subunits of the cnidarian-specific clade evolved from a single precursor. As we have shown here, three of these subunits are activated by Glu. We found that in the three nvpLGICs belonging to clade VII together with the nvGluCls (NV2.181184, NV2.18185, and NV2.19245; Fig. 1), the two critical Lys residues (K74 and K228 in nvGluCl-2) are conserved, strongly suggesting that these receptors are also activated by Glu. However, in most nvpLGIC subunits, these two lysine residues are absent, strongly suggesting that they have another ligand, as we demonstrated for pLGIC-3, pLGIC-4, pLGIC-5, and pLGIC-7. For these four subunits, this ligand is unlikely to be GABA or glycine. The sequence variations in the ligand-binding pockets of the cnidarian pLGICs of the GABA_A_ type suggest the existence of a variety of ligands that are yet to be identified.

We found that nvGluCl-1, nvGluCl-2, and nvGluCl-3 are anion channels like human GABA_A_Rs. This finding is in agreement with the presence of -1′Ala and -2′ Pro residues in M2 (Hibbs and Gouaux 2011). Interestingly, while also nvpLGIC-3, nvpLGIC-4, and nvpLGIC-5 possess -1′Ala and -2′ Pro in M2, nvpLGIC-7 and several other nvpLGICs of the same *Nematostella*-specific clade IV have an Asp residue at position -1′ (Supplementary Fig. 2), which forms a ring of negative charges within the ion pore, strongly suggesting that nvpLGICs of this clade are cation channels. Together, these findings suggest a scenario in which in *Nematostella*, ligand specificity and ion selectivity of pLGICs rapidly diversified starting from a single precursor. We speculate that this specialization of ion channel receptors was instrumental in the evolution of complex behaviours and sensory processing by the cnidarian nervous system (Bosch, et al. 2017). In particular, the diversification of anion channels may have allowed the refinement of inhibitory systems.

Interestingly, lophotrochozoan invertebrates also have inhibitory pLGICs gated by diverse ligands, such as glutamate (GluCls) (Cully, et al. 1994; Cully, et al. 1996; Dent 2006), serotonin (MOD-1) (Ranganathan, et al. 2000) and other biogenic amines (Ringstad, et al. 2009), acetylcholine (ACC, LGC-46) (Putrenko, et al. 2005), or histamine (Zheng, et al. 2002). Thus, it appears that the relatively uniform ionotropic inhibitory signalling with just two ligands (GABA and glycine), which is characteristic of vertebrate nervous systems, is the exception rather than the rule in the animal kingdom.

### Convergent evolution of glutamate-gated ion channels

It is generally believed that glutamate was used as a transmitter for fast ionotropic signalling already at the base of the animal kingdom (Moroz, et al. 2014; Moroz, et al. 2021). As a proteinogenic amino acid it is ubiquitous, there are homologs of vesicular transporters and reuptake transporters, and a large variety of predicted iGluRs in the *N. vectensis* genome (Anctil 2009; Oren, et al. 2014). While formal proof that these receptors bind to and are activated by glutamate is missing, our study establishes glutamate-gated ion channels in the cnidarian pLGIC superfamily. Of note, K74 and K229, which are essential for Glu binding in nvGluCl-2 (Fig. 5), are absent from GluCls in *C. elegans* (Fig. 5a). These GluCls use another Arg residue of loop G to stabilize the α-carboxylates of glutamate (Hibbs and Gouaux 2011). Furthermore, yet another Arg residue in loop A is important for glutamate binding in molluscan GluCls (Fig. 5a), indicating that Glu-binding pockets have evolved independently several times in pLGICs (Lynagh, et al. 2015). Interestingly, while GABA binds to the β+/α– interface of mammalian GABA_A_Rs (Fig. 2a), it has recently been found that glutamate binds to the α+/β– interface to modulate the response of mammalian GABA_A_Rs to GABA (Wen, et al. 2022). Thus, it appears that there is an extensive crosstalk of glutamate with inhibitory GABA_A_Rs across the phylogenetic tree. Some GABA_A_Rs are modulated by Glu, while others (the GluCls) are directly activated by Glu.

### Expression and putative functions of glutamate-gated pLGICs in *Nematostella*

In situ hybridization confirmed that all three investigated nvGluCls are expressed in the tentacles– predominantly in cnidocytes, N2 neurons, or both, and to some extent also in tentacle retractor muscles. Single-cell transcriptome analyses suggest expression of some nvGluCls in specific neuronal subpopulations, which may also be tentacle-specific. The cnidocytes fall into several different types, including nematocytes and spirocytes; spirocytes are restricted to the tentacles. Nematocytes integrate multiple signals and receive synaptic input from spirocytes and sensory neurons (Weir, et al. 2020). Interestingly, while neither Glu nor GABA elicit currents in nematocytes, acetylcholine (ACh) elicits an inward Ca^2+^ current via nAChRs, which leads to the activation of a K^+^ channel and relieves inhibition of voltage-gated Ca^2+^ channels (Ca_v_s), allowing the discharge of the nematocytes (Weir, et al. 2020). In addition, it has been found that the proton-sensitive ion channel NeNaC2 can induce discharge of cnidocytes (Aguilar-Camacho, et al. 2023), illustrating that several transmitters act on cnidocytes. Therefore, we speculate that nvGluCl-2 is expressed in spirocytes, where its activation by glutamate could hyperpolarize the neuron either to release Ca_v_s from inhibition, allowing discharge, or to inhibit discharge. The tentacle retractor muscle is the only ectodermal muscle in *Nematostella* and is one of the two fast-contracting muscles (Cole, et al. 2023). Similar to the discharge of nematocytes, the contraction of the tentacle retractor muscles is regulated by ACh (Faltine-Gonzalez and Layden 2019), and at least some nAChRs are also expressed in tentacles, similar to the nvGluRs in this study. Thus, nvGluCl-2 and, in particular, nvGluCl-3 are likely involved in the regulation of tentacle retractor muscle contraction.

In summary, our study lays the basis for the functional characterization of ion channel receptor diversity in Cnidaria and emphasizes that the overall homology of ion channel receptors does not allow to predict their ligand specificity, which requires detailed structural and functional investigations.

## Methods

### Phylogenetic analysis

Cnidarian pLGICs were retrieved from *N. vectensis* (Zimmermann, et al. 2023) and *H. vulgaris* (Cazet, et al. 2023) proteomes based on the InterproScan annotation as belonging to either the “Ligand-gated ion-channel” or the “Neurotransmitter-gated ion-channel” superfamilies. Human, *Drosophila* and *Caenorhabditis* pLGICs were retrieved from Swiss-Prot (Supplementary Table S1). Datasets per species were deduplicated with CD-hit v4.8.1 (Fu, et al. 2012) using exhaustive search (g = 1) and 99 % sequence similarity (c = 0.99). Deduplicated sequences were aligned using Mafft v7.526 (Katoh, et al. 2002) with the “localpair” algorithm and 1000 optimization iterations (--maxiterate 1000 –localpair). The alignment was trimmed with Trimal v1.5.0 (Capella-Gutierrez, et al. 2009), using the gappyout algorithm (-gappyout). The trimmed alignment was then used for tree construction using Iqtree2 v2.3.5 (Kalyaanamoorthy, et al. 2017) with extended model selection (-m MFP) and 10,000 ultrafast bootstrap calculation (-B 10000) and Baliphy 4.0-1.88.0 (Redelings 2021) using 1 million iterations and summarizing the results using bp-analyze with 150K burn-ins. The iqtree ML tree was selected to represent the topology and node posterior probabilities were exported from the Baliphy 50 % consensus tree. Tree visualization was done using ggtree v3.12.0 (Xu, et al. 2022).

### Molecular biology

Full-length DNA sequences of GABA_A_ subunits were identified and cloned based on bulk and single cell transcriptomes of *N. vectensis* (Steger, et al. 2022; Cole, et al. 2023; Cole, et al. 2024). nvpLGIC-1 – nvpLGIC-4 were cloned in oocyte expression vector pRSSP6009 (Bässler, et al. 2001). nvpLGIC-5 and nvpLGIC-7 DNA sequences were synthesized and cloned in the pCDNA3.1 (-) vector (BioCat GmbH). They were then subcloned in the vector pRSSP6009 (nvpLGIC-6) or pRSSP 6013 (nvpLGIC-5). Site-directed mutations were introduced in nvGluCl-2 to create the mutants K229T, using the Quick-change mutagenesis method. nvpLGIC-1-nvpLGIC-6 subunits were fused with His_6_-tags at their C-terminus using PCR. His_6_-tags were incorporated just before the stop codon without changing any other amino acids. All final constructs were sequenced for confirmation. The plasmid containing full-length cDNA for human GABRB3 was purchased from the Harvard Plasmid Repository (order number HsCD00043103; repository closed in January 2021) and subcloned into a Gateway-compatible version of the pNKS2 vector (Gloor, et al. 1995) using the Gateway cloning system (Invitrogen, Karlsruhe, Germany). cRNA was synthesized using SP6 RNA polymerase from linearized cDNA using the mMessage Machine kit (Ambion).

### Blue native gels and electrophysiology

Approximately 2-5 ng of cRNA of nvpLGICs were injected into *Xenopus laevis* oocytes of stages V and VI. Injected oocytes were incubated at 19° C in oocyte Ringer’s solution (OR-2) containing (in mM): 82.5 NaCl, 2.5 KCl, 1.0 Na_2_HPO_4_,1.0 MgCl_2_, 1.0 CaCl_2_, 5.0 HEPES, 0.5 g/l PVP, 1,000 U/l penicillin, and 10 mg/l streptomycin. pH was adjusted to 7.3 using NaOH.

For blue-native gels, His-tagged nvpLGIC-1-nvpLGIC-3, nvpLGIC-5, and human GABA_A_ β3 were expressed in oocytes, purified and resolved by SDS-PAGE and blue native (BN) PAGE as described (Nicke, et al. 1998; Haeger, et al. 2010). In brief, defolliculated, cRNA-injected *X. laevis* oocytes were metabolically labelled with [^35^S]methionine, and, after 2 days, surface-labelled with the membrane-impermeant dye IR800 NHS ester (LI-COR) for 1h. After digitonin solubilization, receptors were purified via their C-terminal His tag using Ni-NTA-Sepharose and resolved by BN-PAGE in their native and partially LiDS-denatured state to visualize their oligomeric state. IR800 fluorescence was detected on the wet BN-PAGE gel using a LI-COR Odyssey scanner, while ^35^S was detected after drying of the same gel using phosphorimaging. Protein expression was displayed using Image Lab (Bio-Rad).

For electrophysiology, whole-oocyte currents were measured two days after injection using a two-electrode voltage clamp (TEVC) set-up with a TurboTec 03X amplifier (npi electronic GmbH, Tamm, Germany) and a pump-driven solution exchange system combined with the oocyte testing carousel (OTC), controlled by the interface OTC-20. For nvGluCl-1, the His-tagged version was functionally characterized. Standard bath solution contained (in mM): 140 NaCl, 10 HEPES, 1.8 CaCl_2_, and 1.0 MgCl_2_, pH was adjusted to 7.4 using NaOH. Solution exchange was controlled using the CellWorks 5.1.1 software (npi electronic). Data were sampled at 0.1 - 1 kHz and filtered at 20 Hz. Holding potential was -70 mV.

### Homology modelling and Molecular docking

The homology model of pLGIC-2 was designed using the SWISS-MODEL server (Waterhouse, et al. 2018) using the crystal structure of the human β3 homopentamer (PDB ID: 4COF) (Miller and Aricescu 2014). The pLGIC-2 monomer was aligned and superimposed onto each of the five monomers of the homopentameric model using PyMOL (The PyMOL Molecular Graphics System, Version 2.5.4, Schrödinger, LLC), resulting in a complete pLGIC-2 homopentamer model superimposed onto the 4COF template. The alignment resulted in a root mean square deviation of 0.16 Å between the Cα atoms of the modeled chain and the template.

A model of the GABA-pLGIC-2 liganded receptor was studied using molecular docking simulations. Receptor and ligand structures were prepared for docking using the open-source AutoDock Tools 1.5.7 (Morris, et al. 2009) and binding affinities and optimal ligand-receptor binding poses were predicted using AutoDock Vina (Trott and Olson 2010). For this calculation, a 60 × 56 × 56 Å³ box was constructed around the GABA-binding pocket, approximately centered on the Cα atom of F93. The algorithm’s exhaustiveness was set to 8, with an energy range of 4. Molecular interactions between the receptor and ligands were analysed using LIGPLOT (Wallace, et al. 1995) and the molecular docking results were visualized with Chimera X 1.7.1 (Goddard, et al. 2018) to inspect their interaction in detail.

### Single cell analysis

Expression profiles of all genes of interest were examined across cell state clusters described in(Cole, et al. 2024). The published dataset was imported to Seurat Vs.4 and expression profiles were visualized with the Seurat::DotPlot function. Cell-states with above average expression in at least 20% of the cells were deemed positive for expression of the putative receptor genes.

### Fluorescent and colorimetric *in situ* hybridizations

For whole-mount *in situ* hybridization (ISH) embryos were fixed 0.25% glutaraldehyde/3.7% formaldehyde/ PTW (PBS + 0.1% Tween), then in 3.7% formaldehyde/PTW for 1 hr at 4°C. Fixed embryos were washed 4x with PTW and stored in methanol at -20°C. Colorimetric ISH was performed as previously described (Kraus, et al. 2016) with minor adaptions: samples were incubated in anti-Digoxigenin-AP Fab fragments (Roche) diluted 1:4000 in 0.5% blocking reagent (Roche)/1x MAB (Roche) ON at 4°C. After 10x 10 min washes in PTW, samples were rinsed 3x 5 min with alkaline phosphatase buffer and stained with 1µL NBT and 1.5 µL BICP in 1 mL alkaline phosphatase buffer as described previously (Kraus, et al. 2016). Embryos were embedded in 86% glycerol and imaged with a Nikon 80i compound microscope equipped with a Nikon DS-Fi1 camera.

Fluorescent ISH was performed according to Tournier et al. (Tourniere, et al. 2020) with minor adaptations in antibody incubation: Samples were blocked in 0.5% blocking reagent (Roche)/Maleic acid buffer (Roche) for 1 hr at room temperature. Anti-Digoxigenin-POD Fab fragments (Roche, 11633716001) and anti-fluorescein-POD Fab fragments (Roche, 11426346910) were diluted 0.15 U/mL in blocking solution. After overnight incubation, samples were washed 10x 10 min in TNT (0.1M Tris-HCl pH 7.5/0.15M NaCl/0.5% Triton X-100), and then incubated in fluorophore tyramide amplification reagent (TSA® Plus Kit, PerkinElmer) for 30 min at room temperature. Following the staining reaction, samples were washed with TNT. For double staining samples were incubated in 2% H_2_O_2_/TNT for 20 min at room temperature after TNT washes and then incubated again in blocking solution before adding the second antibody. Finaly samples were mounted in VECTASHIELD® antifade mounting medium (Vector Laboratories) and imaged with a Leica STELLARIS 5 confocal microscope. Maximum intensity projections were generated using Fiji (Schindelin, et al. 2012).

### Statistical analysis

*Xenopus* oocytes were randomly assigned to experimental groups without blinding of the experimenter. For electrophysiological experiments, we used cells from a minimum of two batches of oocytes isolated on different days from distinct animals. Each oocyte served as a biological replicate for electrophysiology.

Electrophysiological data were analysed offline, and current amplitudes were assessed using CellWorks Reader 6.2.2 (npi electronic GmbH, Tamm, Germany). Concentration-response relationships were normalized to the currents obtained with a saturating concentration (*I/I_max_*). The mean of normalized amplitudes was plotted as a function of ligand concentrations. EC_50_ values and Hill coefficients H were calculated by fitting the data to the Hill equation:

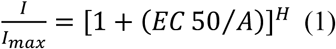

using IGOR Pro (version 8.04, waveMetrics, Inc.), where *I* is the current amplitude, *A* is the concentration of the ligand, EC50 is the concentration of the ligand that produces half-maximal effect, and *H* is the Hill coefficient.

Desensitization kinetics and recovery from desensitization were fitted to a mono-exponential function:

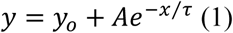

where y_o_ is the offset, A is the amplitude and τ is the time constant. A fit with a double exponential function did not yield substantially different results, because > 90 % of the current decline was described by one component, which was similar to the time constant obtained with a single exponential fit.

To calculate the reversal potential, mean current amplitudes were plotted as a function of voltage and were fitted using a linear equation. The voltage, at which the direction of current reversed from negative to positive, was considered as reversal potential. Data are presented as mean ± standard deviation.

## Data Availability

Datasets generated and analysed during this study are included in this published article. Single cell data is available from the UCSC Cellbrowser at “sea-anemone-atlas.cells.ucsc.edu”. Additional information is available from the corresponding authors upon reasonable request.

## Acknowledgments

Confocal microscopy was performed at the Core Facility Cell Imaging and Ultrastructure Research, University of Vienna-member of the Vienna Life-Science Instruments (VLSI). This work was supported by a grant of the Deutsche Forschungsgemeinschaft (GR1771/8-1) to S.G. and by the ERC-AdG EvoNEUROMUSCLE to U.T. We thank Adrienne Oslender-Bujotzek for excellent technical assistance.

## Author contributions

AO and SJ performed the functional characterization by two-electrode voltage clamp; MB and SA performed homology modelling and AOR performed molecular docking and visualized the model; MR performed blue native PAGE; AGC carried and analysed out single cell RNAseq analyses, JM performed the phylogenetic analyses, SK and LH cloned the LGIC1-7, LK performed in situ hybridisation experiments, UT and SG designed the study; GS, UT, and SG supervised the study; SG wrote the original draft and all other authors reviewed and edited the original draft.

