## Supplementary information for "Functional analysis of ligand-gated chloride channels in a cnidarian sheds light on the evolution of inhibitory signalling"

**This Supplement includes:**

Supplementary Figs. S1 to S4

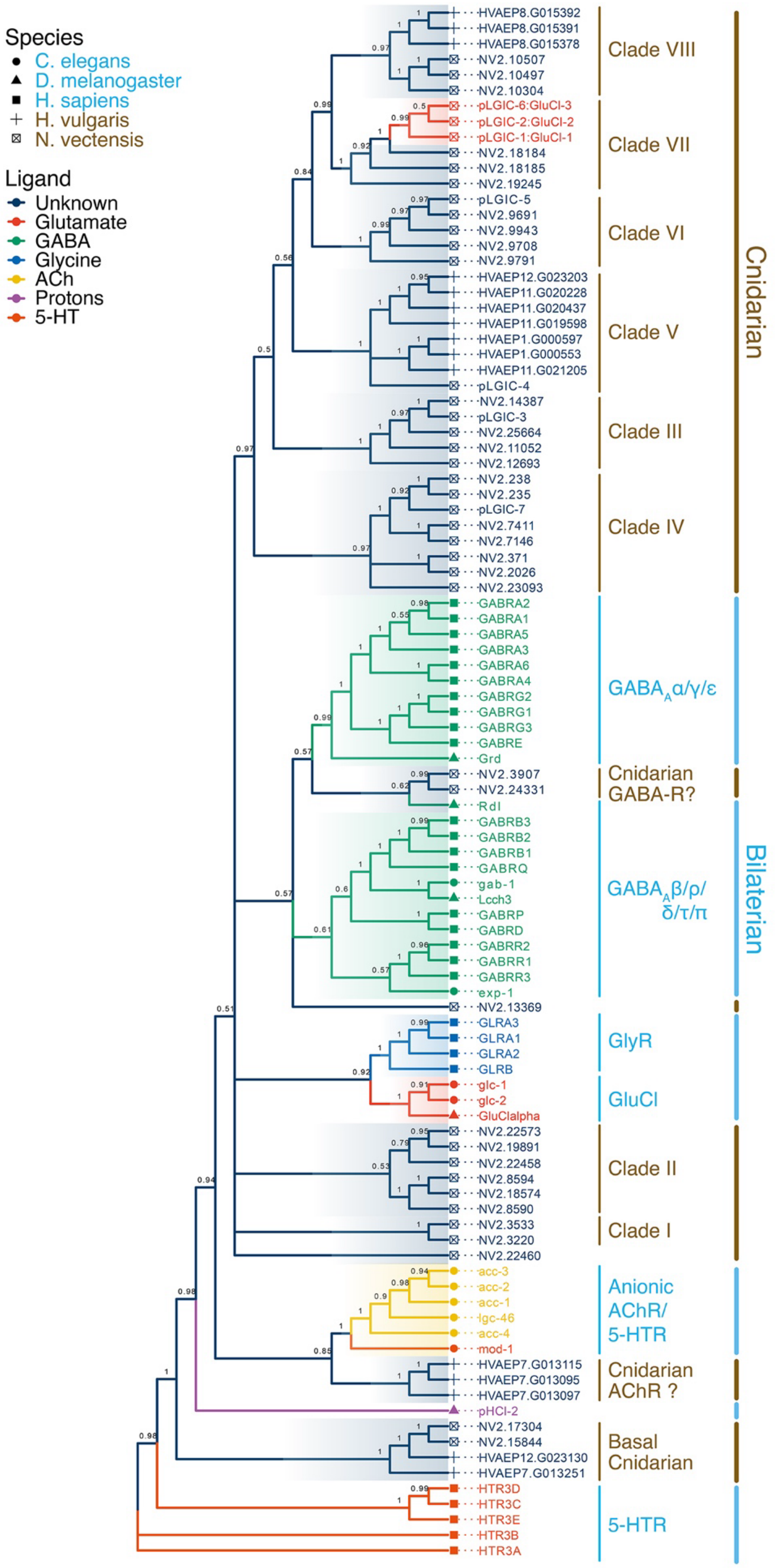


**Supplementary Fig. S1. Phylogenetic analysis of cnidarian $GABA_A$Rs/GlyRs.** Bayesian analysis of pLGIC genes in *N. vectensis*, *H. vulgaris*, *C. elegans*, *D. melanogaster*, and *H. sapiens*. Annotation of two receptors from *N. vectensis* as "Cnidarian GABA-Rs" and of three from *H. vulgaris* as "Cnidarian AChR" is provisional as it has relatively low support. Posterior probabilities for individual nodes are indicated. The tree was rooted with mammalian $5\text{-}HT_3$Rs as an outgroup.

```
Hs-GBRA1      -----------MRKSPGLSDCLWAWILLLSTLTGRSYGQPSLQDELKDNTTVFTRILDRLL  50
Hs-GBRB3      --------------MWGLAGGRLFGIFSAPVLVAVVCCAQSVNDPG--NMSFVKETVDKLL  45
nvpLGIC-1     ----MRAFWSGCVCVLTAAATAAQSLSNSDESSRTERSFDEYYPEMEEARNLSDFINNAL  56
nvpLGIC-2     ----------MYIAAVYVAICVFLMGCISANCSSWPEDQRKDEGNPIANAHNLSVFINDAL  51
nvpLGIC-3     MACRMSKIRYFLQEWIFKGD-KMKLILLLGLFYMVECYSNPSPNSSNQTVSRILDSILTA  59
nvpLGIC-4     ----------MELALWLFVMATLMTMSIWQSEAQFSQSTFSGKSKDDIATLTATKRLHSML  51
nvpLGIC-5     ---------------------MEVIASILLLGLSLQVNANDDFN--------TSKAIDKLFSK  34
nvpLGIC-6     ------------MSWRRVWLALFLGCLSLLFCCAEEKEITDEMRKSMEEARNLSLFINNVI  49
nvpLGIC-7     --------------------------MILSREANFIWKFLLLSLFTRHAQCSMEGPVLKMLF  36

                                       G                   D
Hs-GBRA1      DGYDNRLRPG-LGERVTEVKTDIFVTSFGPVSDHDMEYTIDVFFRQSWKDERLKFKGPMT 109
Hs-GBRB3      KGYDIRLRPD-FGGPPVCVGMNIDIASIDMVSEVNMDYTLTMYFQQYWRDKRLAYSGIPL 104
nvpLGIC-1     KIHDKKVRPN-AGGPPVMVQVEFKVVSIGEIKEAEMVYSMDIFFRQWWYDPRFAHNFSKP 115
nvpLGIC-2     ASYDRKVRPN-YGGLPVMVLIEFKVISFGEIQEANMEYSFDIFLRQWWYDPRFKSNFTEP 110
nvpLGIC-3     EKYDKRLRPN-YGAGPVIVTVGFWVLSIDTINVVDMDYTLDIFLRQSWVDERLSHDLNTT 118
nvpLGIC-4     EGYESKIRPN-YQGDPVEILIDISVASFGNLQEADMKFSLDMFFRQSWHDPRLRHNMNET 110
nvpLGIC-5     ESYDRRLRPHRQTGRLINVTIGMTVVSFGQIRERDMDFSLDIFFYQKWSDPRLKHSLKDP  94
nvpLGIC-6     RRHPRNVRPN-AGGEKVMVDIEFKVISFGEIKEANMEYSLDIFFRQWWYDSRFAHNYSMP 108
nvpLGIC-7     DGYDKEARPN-WGGRPVEVRTDAIVEAFGNIQEANMEYPVQLYFHQYWTDKRLAGKLNLS  95

                                A                            E
Hs-GBRA1      VLRLNN--LMASKIWTPDTFFHNGKKSVAHNMTM-PNKLLRITEDGTLLYTMRLTVRAEC 166
Hs-GBRB3      NLTLDN--RVADQLWVPDTYFLNDKKSFVHGVTV-KNRMIRLHPDGTVLYGLRITTTAAC 161
nvpLGIC-1     FTMAA---DATQLFWTPDTYFWNVKNAKYHRVTR-ENMRVMINPDGKIYFSTRITITAQC 171
nvpLGIC-2     FTMAA---DPTKMFWTPDTYFWNVKRSKYHHVTR-ENMRVMIWPDGKIYFSTRITLTAQC 166
nvpLGIC-3     MFLSN---TVMDKIWLPDSYFVNAKQGSFHKVTK-DNMMVMIQPDGQVQYNARVTIRASC 174
nvpLGIC-4     MTLTTGTKHPADFVWVPDTVFVDASHAYMHNVMV-ANHKLDVTPGGRVFWGTRATLIARC 169
nvpLGIC-5     IILGG---EFKKLVWLPDTFFLNIKTAKFHTVPS-DNSKISIFEDGTVRYSTRLTLTAGC 150
nvpLGIC-6     FTMAA---DPTELFWTPDTYFWNVKSANYHRVTR-ENMRVMINPDGKIYFSARITLTCQC 164
nvpLGIC-7     ITLTVS---AASNACKPDPYCYNARQSNMMLEDKDVNSRLSIDPDGKIFYSRGVSIVASC 152

                   Cys-loop             B                     F
Hs-GBRA1      PMHLEDFPMDAHACPLKFGSYAYTRAEVVYEWTREPARSVVVA-EDGSRLNQYDLLGQTV 225
Hs-GBRB3      MMDLRRYPLDEQNCTLEIESYGYTTDDIEFYW-RGGDKAVTGV--ERIELPQFSIVEHRL 218
nvpLGIC-1     DMDLRLYPMDIQYCPLIIESYAHTRSDVDYTW-KGG--DDQGVEIVSKEMAQFEFIGANT 228
nvpLGIC-2     DMDLRLYPMDTQHCPLIIESYAYTKNDLDYKW-------NSQGIEIVSSEMAQFDLIKVHT 220
nvpLGIC-3     PMNLRKFPMDTQHCPLTLESYGYSSDHIVFKWEIEDGDGLGFVPESLKMLPQYKLARVHL 234
nvpLGIC-4     HMNLRRYPMDEQTCNLSIISYAYPVRHLLYRW-K-----NSPGIKVLDGEMSQFYMTRITT 224
nvpLGIC-5     EMNLLDYPLDEQTCNLTILSYAFSTHEMDYIW-DGG--TKSAIDVINDAMNEFTLIGINT 207
nvpLGIC-6     EMDLHLYPLDTQECPLRIESYAHTVADVDYRW-KGG--ETQGVEIVSEEMAQFDLLGVRT 221
nvpLGIC-7     EMDLHDFPLDTQNCYLKFGSYAFTDTDIFFRW----NNPNKLSRVQKSDLAQFHLAGYDL 208

                     C                         M1
Hs-GBRA1      DSGIVQSSTGEYVVMTTHFHLKRKIGYFVIQTYLPCIMTVILSQVSFWLNRESVPARTVF 285
Hs-GBRB3      VSRNVVFATGAYPRLSLSFRLKRNIGYFILQTYMPSILITILSWVSFWINYDASAARVAL 278
nvpLGIC-1     YTKAQTNSKGSFASLRAEFTFKRRVAYFITATYMPAMILVILSWCTFWIHRNAVPARVTL 288
nvpLGIC-2     TSKKQENSKGSFASLMAIFSFRRRTESFVSSIYVPSVVLVVLSWCCFFINPDAVPARVGL 280
nvpLGIC-3     STLHNEYVVGNWSGIKALFTFERMYSYFVIHVYGPCALIVVVSWVSFLLPKEQAPARITL 294
nvpLGIC-4     SLRNEIYVAGEYSVLKASFTFQRRVGFYLIQIYIPCMAIVFVAWMSLWVDRRATPARVSL 284
nvpLGIC-5     SKQVFRYVTGPWTHLEATFKFKRRLGYSIIQVYAPTILIVALSWLSFWISKEAAPARVAL 267
nvpLGIC-6     DTKKSTNSKGAFASLKATFRFRRRMDFYLSSIYVPEVILVVLSWTTFFITPSAVPARTAL 281
nvpLGIC-7     ITETGIYEEGNYTTVKVIFKMNRRVGYYIIQAYCPDVLIVVLSWIIFWMDVKDMGDRMAL 268
```

```
                 M2                              M3
Hs-GBRA1    GVTTVLTMTTLSISARNSLPKVAYATAMDWFIAVCYAFVFSALIEFATVNYFTKRGYAWD 345
Hs-GBRB3    GITTVLTMTTINTHLRETLPKIPYVKAIDMYLMGCFVFVFLALLEYAFVNYIFFGRGPQR 338
nvpLGIC-1   SITTILTTILLTSSTNGSMPKVSYSKAIDYFLMTSLGFIFMSLLEYILVLNT----HPNF 344
nvpLGIC-2   SITTILTSILLQGSVNSNMPKVSYMKAVDYFLLTSFGFIFAALLEYIMVLNT----DANL 336
nvpLGIC-3   GVTSVLTVVTILNMLNNSMPKVNYVKTIDKYLIGCFLFVFATLVEYSLVLWLTKSYKKYN 354
nvpLGIC-4   CITTLLTIATIWGNVNASMPRVSYVKSIDIYLMTSFTFVLGTLLEYIVIMNKGHPKSKKI 344
nvpLGIC-5   GITTVLTIVTLMGSFRAAVPKVSYVKAIDLFFIVSFFFVFGAVMEYVAVLLHSAMVDANK 327
nvpLGIC-6   SITTILTTILLSSSVNSGMPKVSYMKSIDHFMLISFGFIFAALIEYIIVLNS----PSRF 337
nvpLGIC-7   GITTILTIMFLLGAVNASMPKVSYPKALDWYLMVSFAFVFLTLIESMIVFLLTPQGESEK 328

Hs-GBRA1    GKSVVPEKPKKVKDPLIKKNN------TYAPTATSYTPNLARGDPGLATIAKSATIEPKE 399
Hs-GBRB3    QKKLAEKTAKAKNDRSKSESNRVDAHGNILLTSLEVHNEMNEVSGGIGDTRNSAISFDNS 398
nvpLGIC-1   WKEKDNVDAEIGLNSVGKKLSPGKDASPEVIVTMMDKKGEESSCKPCSVQT--------- 395
nvpLGIC-2   G---YTRCKMLRHKEQADLAMEDEYAADVDIVVKVMDEGVCRIRNKKGSIK--------- 384
nvpLGIC-3   KAKEELSKSRG-KSHPFDEQYDGDRNTQIALMDMPHENDVVVEMNGRHAAPRHRKIRCQS 413
nvpLGIC-4   RKSVHLNDYMSPYTYSFRNWPDSQTSTTASNINQKNSNTCSKQPEVKDSEP--------- 395
nvpLGIC-5   KSQDNGEMKDVEMQELVKQDPEKVTKQNGDKDNPASPSPSNESLRKRKSTP--------- 378
nvpLGIC-6   T---ECFPMLFRTYSIPEEAQLHKSDGSIANRKPAEPS---------------------- 372
nvpLGIC-7   PKKTSCLTKRVVTMLTPVLRRRPNSHVVTTPNNHELNNISPTDAALLNPEDGGGREGPNG 388

                                                                     M4
Hs-GBRA1    VKPETKPPEPKK------------------------------TFNSVSKIDRLSRIAFPL 429
Hs-GBRB3    GIQYRKQSMPREGHGRFLGDRSLPHKKTHLRRRSSQLKIKIPDLTDVNAIDRWSRIVFPF 458
nvpLGIC-1   --CIAPPPPKPKKPND--------------------------LPYLDVHWSTAPRGFFPS 430
nvpLGIC-2   --VQAEATKVSCRSNR--------------------------LIRRDHWIDRVSRIIFPS 416
nvpLGIC-3   PQEPSIRCEPIKRTLL--------------------------NEAFVQHVDEYALVLFPG 447
nvpLGIC-4   -----------------------------------------------LKVEVYARIGFLS 408
nvpLGIC-5   --ESSCKPRTVVRFAF--------------------------IEHNADIIDRLSRVLFPL 410
nvpLGIC-6   ---------------------------------------------KTHWIDRFSRYFFPI 387
nvpLGIC-7   TKACWDDHTGHVKESK----------------------------CAGYRVDQISRVLFPL 420

Hs-GBRA1    LFGIFNLVYWATYLNREPQLKAPTPHQ                                  456
Hs-GBRB3    TFSLFNLVYWLYYVN                                              473
nvpLGIC-1   STSSSLLLTGSTTSTIPRRTSNSFELRFLDSSRLRLHRLPLNQPQEHPNG           480
nvpLGIC-2   AYVIFIICYTAYYVNKRDNSGEDPPVTTKPASRT                           450
nvpLGIC-3   SFVIFNAVYWFSVMSFDQIFGSFGAIFTLSAHPVCAPVPLTSS                  490
nvpLGIC-4   AFVLFNGVYWAVLLI                                              423
nvpLGIC-5   SYLIFNIIYWMYYTLSSFNS                                         430
nvpLGIC-6   SYSIFFLAYWIHYQSQRQ                                           405
nvpLGIC-7   AFVFYNVAYWYTYLERIPVLGESDIM                                   446
```

**Supplementary Fig. S2.** Sequence alignment of nvpLGICs with human $GABA_A$ α1 and β3. Conserved residues are indicated by white letters on black background. Residues important for ligand binding, ion selectivity and assembly as homomers are indicated by colored letters. Bars represent loops A-G and the transmembrane helices M1-M4; the blue bar represents the Cys-loop. The predicted signal peptides of nvpLGICs are shown in grey; they have been predicted using SignalP 6.0 (Teufel, et al. 2022).

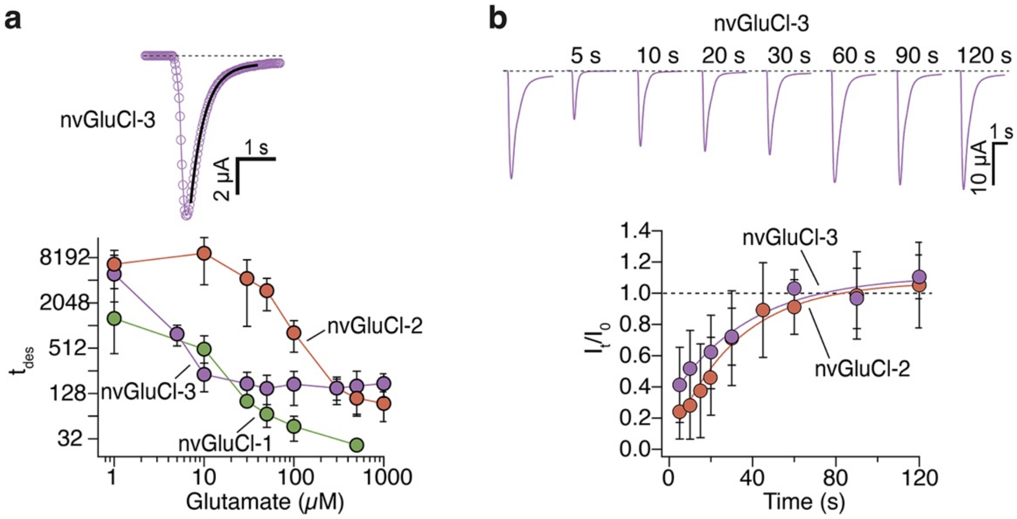


**Supplementary Fig. S3. a)** Top, representative current trace of nvGluCl-3 currents evoked by 100 µM glutamate. Current decline was fitted with a mono-exponential function (black line) to estimate $\tau_{des}$. Bottom, mean $\tau_{des}$ at increasing concentrations of glutamate for nvGluCl-1 (green), nvGluCl-2 (orange), and nvGluCl-3 (violet), respectively. **b)** Top, representative current traces showing recovery of nvGluCl-3 from desensitization induced by application of 30 µM glutamate for 30 s. Currents after washout of glutamate for 5, 10, 20, 30, 60, 90, and 120 s are shown. Bottom, mean normalized current responses of nvGluCl-2 (orange) and nvGluCl-3 (violet) as a function of time. Lines represent fits to a mono-exponential function. For nvGluCl-2, a 30 s desensitizing pulse of 100 µM glutamate was used. Data represent the mean ± s.d. of 7-22 cells.

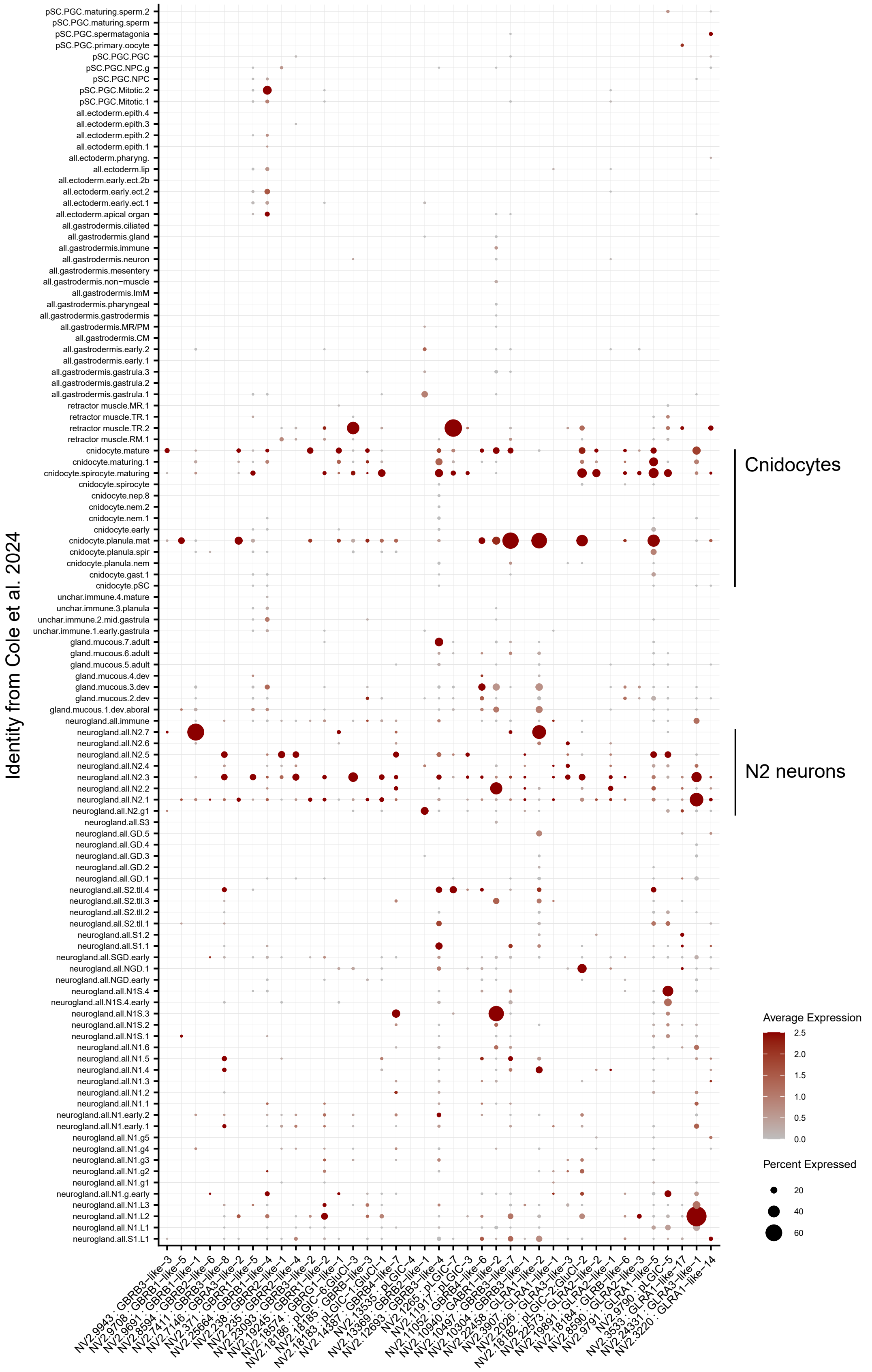


**Supplementary Fig. S4.** Expression profiles of all putative GABA$_A$R-like receptors on the single cell dataset from (Cole, et al. 2024). Only gene models with at least three reads in the dataset are shown. Most models are expressed at low levels and are concentrated primarily within the mature cnidocytes and class N2 neurons ("neurogland.all.N2").
